# TileScope quantifies spatial gene expression relationships beyond colocalization

**DOI:** 10.64898/2026.10.29.762851

**Authors:** Maialen Arrieta-Lobo, Sébastian Lillo, Benjamin Dartigues, Domitille Chalopin-Fillot, Maya Saleh, Thomas Daubon, Macha Nikolski

## Abstract

Gene expression patterns within the same tissue region can exhibit similar correlation or enrichment, but still differ in their organization within regions and at their boundaries. We present TileScope, a framework for analysing genes and transcriptional programs within predefined tissue regions, their components and their contacts with neighbouring regions. TileScope combines regional enrichment, within-region correlation, cross-correlation over distance and expression differences near specific boundaries. Repeated spatial partitions and consensus aggregation show how these measurements depend on the grouping of spots or cells. Across 30 controlled constructions derived from glioblastoma and hepatocellular carcinoma data representing ten spatial expression scenarios, combined measurements separated arrangements that individual statistics left ambiguous, including coincident patches, gradients and expression localised at region boundaries. In 30 human glioblastoma sections, TileScope distinguished shared regional enrichment from internal covariation and identified hypoxia-associated expression localised at region boundary despite negligible within-region correlation. In newly acquired Visium profiles of control and myeloid-targeted MCT1-knockout (KO) orthotopic GL261 mouse gliomas, proliferation and myeloid programs were enriched in tumour peripheries but showed opposite within-periphery-region correlations. Proliferation was higher near the tumour-facing boundary in both samples, whereas myeloid expression was higher near the tumour-facing boundary in the control and farther within periphery in the KO condition. Xenium applications on glioblastoma and breast cancer samples demonstrated feasibility with irregularly spaced individual cells and different regional annotations. TileScope provides spatially explicit descriptions of gene and program relationships, linking their organization within tissue compartments to their sensitivity to aggregation.

## INTRODUCTION

Spatial transcriptomics maps gene expression within tissues, linking expression patterns to anatomical structure (Ståhl et al. 2016; Janesick et al. 2023). The biological information in these datasets extends beyond the abundance of individual genes or cell populations. Mixed and compartmentalised tumour- immune architectures have been associated with outcome in breast cancer (Keren et al. 2018); coordinated expression programs form distinct immune hubs in colorectal cancer (Pelka et al. 2021); and glioblastoma cell states and microenvironmental programs exhibit a layered organization associated with hypoxia (Greenwald et al. 2024). In each setting, where expression occurs and how it is arranged relative to neighbouring tissue are important to its interpretation.

Interpreting these relationships requires more than identifying a pair of genes or programs as enriched, colocalised or correlated. Two programs can be enriched in the same region yet vary independently within it, or be strongest in opposing parts of that region. Conversely, positive correlation can arise from broad overlapping expression, coincident foci or gradients in the same direction. A program may also be elevated near a particular neighbouring compartment without correlating with another program across the entire region. The same pair can therefore show different spatial relationships in different parts of a tissue, while several arrangements can coexist within one annotated region.

Computational methods address distinct aspects of these relationships. MERINGUE uses spatial autocorrelation and cross-correlation to identify spatial expression patterns and relationships between genes (Miller et al. 2021). SpatialCorr and spCorr estimate or test spatial variation in gene correlation, including differences within or between tissue regions (Bernstein et al. 2022; Jiang et al. 2026). SpaceBF instead models one gene’s expression as a spatially varying function of the other, estimating local regression coefficients with shrinkage across neighbouring locations (Seal and Neelon 2026), while STopover identifies connected components in expression maps and quantifies their overlap (Bae et al. 2025). These approaches quantify different aspects of spatial expression. However, a correlation calculated across all spots or cells in a tissue region does not show whether two expression patterns vary together throughout that region or only within a few local patches. Expression may also be concentrated near a neighbouring compartment despite weak correlation across the region. A relevant question that requires systematic solution is where their expression overlaps or separates and how those patterns relate to the surrounding tissue.

Spatial analyses often combine nearby measurements to reduce sampling noise or study organization across groups of cells. For example, (Greenwald et al. 2024) quantified cell state composition within sliding windows of different sizes to examine relationships beyond immediate neighbours. geneSCOPE pools transcripts into spatial bins before estimating gene–gene relationships, selecting a bin size that balances statistical stability and spatial detail (Zhang et al. 2026). These summaries can reveal broader patterns, but larger units can also blur boundaries and merge distinct expression patterns. Correlation estimates may change with both the size of the units and the way observations are assigned to them. This dependence is known as the modifiable areal unit problem (MAUP; Openshaw 1984; Lloyd 2014; Manley 2014; Stillwell et al. 2018). Comparing alternative groupings of the same observations makes it possible to assess how much a reported relationship depends on the chosen partition.

Here we present TileScope, a framework for analysing spatial relationships between genes or transcriptional programs within predefined regions of interest (ROIs) and at their boundaries. It distinguishes enrichment from internal covariation within ROIs, association over distance and expression near a neighbouring region, representing disconnected ROI components and contacts explicitly. Repeated tessellations vary how spots or cells are grouped into tiles, revealing sensitivity to the placement of partition boundaries, the zoning effect of the MAUP. Comparing statistics calculated at individual spots or cells with those calculated on the consensus of these partitions shows how expression relationships change after aggregation. We evaluated these measurements on controlled spatial scenarios built on tissue geometries observed in experimental Visium 10x Genomics data comprising eight human glioma samples from Greenwald et al. (2024), three human hepatocellular carcinoma samples from Giraud et al. (2024) and two out of the three orthotopic GL261 mouse glioma samples acquired for the present study. In human glioblastoma, combining enrichment, correlation and boundary measurements allowed TileScope to distinguish local arrangements underlying the program relationships described by Greenwald et al. (2024). TileScope showed that the same pair of programs could be enriched in two ROIs but vary together only in one. It also identified higher hypoxia-associated mesenchymal program scores near an ROI boundary, identifying a pattern that the ROI- wide correlation failed to capture. We then applied the same analytical approach to Visium profiles of orthotopic GL261 mouse gliomas generated for this study. Enrichment identified peripheral tumour regions containing both proliferation and myeloid programs in samples from control mice or mice with a myeloid- targeted deletion of the monocarboxylate transporter (MCT1, encoded by the *Slc16a1* gene), dubbed as *Mct1^ΔMye^* mice (or knockout [KO] herein). Correlation and boundary measurements together distinguished the organization of the programs in the peripheral tumor region: while the proliferation and myeloid programs varied together near the tumor-facing boundary in controls, myeloid-targeted deletion of MCT1 induced a distinct spatial segregation. Within the tumor periphery of KO samples, proliferation scores were highest near the tumor-facing boundary, whereas myeloid scores peaked at the outermost boundary. The combined statistics thus distinguished aligned from opposed expression patterns within similarly enriched tumour compartments and located those patterns relative to tumour anatomy. Applications to publicly available glioblastoma and breast cancer Xenium datasets (Meylan et al. 2026; Janesick et al. 2023) further demonstrated feasibility at individual cell resolution.

Together, these analyses show how regional context, distance and boundary position distinguish arrangements that enrichment or correlation alone leave unresolved, while making sensitivity to aggregation explicit.

## RESULTS

### Overview of TileScope

TileScope analyses pairs of continuous expression features, such as genes or transcriptional program scores, within predefined regions of interest in spatial transcriptomic data (Fig. 1). Inputs comprise expression values, observation coordinates and ROI labels, with observations corresponding to either spots or individual cells. ROIs are user-defined, either by expert knowledge (e.g. anatomical annotation) or by data- driven approaches such as clustering algorithms like BayesSpace (Zhao et al. 2021). The framework preserves these biological region labels while distinguishing their spatially disconnected components and boundaries with neighbouring ROIs. This distinction allows regional expression relationships, organization within individual components and expression near particular boundaries to be examined without conflating their spatial contexts. Application to Xenium samples demonstrated feasibility on irregularly distributed individual cells, with regional statistics and contributing cell counts examined before and after aggregation (Supplementary Fig. S7-S8; Supplementary Section 6; Supplementary Data 3).

**Figure 1.**
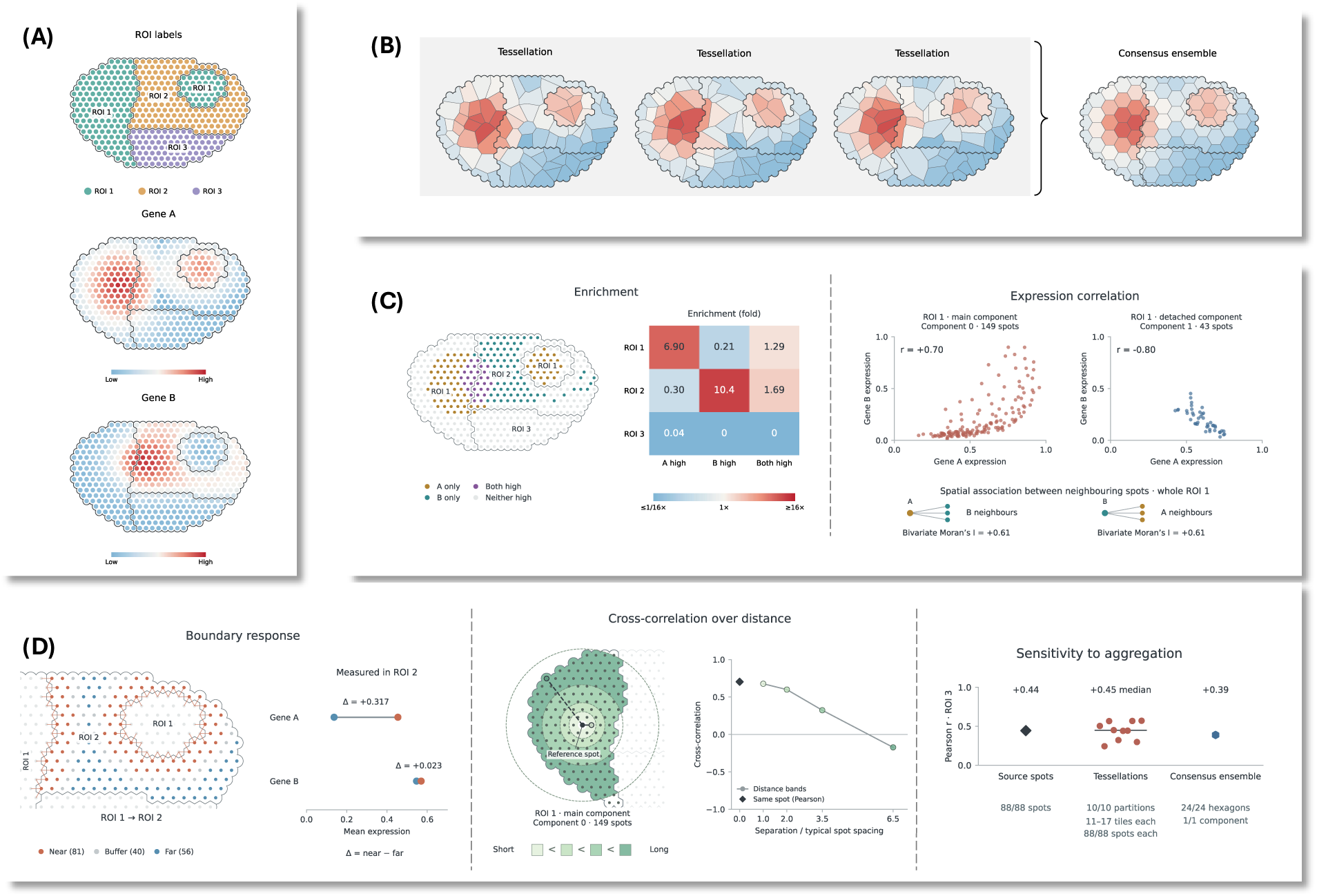
Overview of TileScope. **(A)** Synthetic tissue with three ROIs and expression of two genes. ROI 1 contains two disconnected components. Blue to red indicates increasing expression. **(B)** Three Voronoi tessellations of the same synthetic tissue illustrate how Gene A expression from (A) is averaged within tiles, projected onto a hexagonal grid and combined into an ensemble consensus Gene B is treated identically, using the same partitions. **(C)** The map identifies spots with high expression of either gene or jointly of both genes; enrichment compares their prevalence within each ROI with the other ROIs. Scatterplots show contrasting Pearson correlations in the two ROI 1 components. Bivariate Moran’s I relates Gene A expression at each spot to Gene B expression in neighbouring spots, and vice versa, across ROI 1. **(D)** From left to right: boundary response, expressed as Δ = (mean expression near the boundary – mean expression far from the boundary), calculated separately for each gene within ROI 2 (responder ROI) at its contacts with ROI 1 (anchor ROI); spatial cross-correlation between Gene A and Gene B across four distance bands within the main ROI 1 component, concentric circles illustrate the distance bands around one reference spot and the cross-correlation curve summarizes comparisons across the component; and sensitivity of ROI 3 correlation to aggregation, comparing individual spots, ten tessellations and ensemble consensus, with contributing support indicated.

TileScope repeatedly partitions each connected component into tiles and averages gene expression values or program scores within each tile. It compares relationships measured at individual spots or cells with those obtained from these alternative partitions and from a common hexagonal consensus grid. Different tessellations provide alternative spatial groupings of the same spots or cells. Expression summaries from these partitions are combined on a common hexagonal consensus grid using the overlap between tiles and hexagons, following the principle of areal interpolation used in geographical analysis (Goodchild and Lam, 1980). Together, these representations allow relationships to be examined before and after aggregation, rather than relying on a single partition of the tissue.

The statistical outputs address complementary questions about spatial organization. Marginal and joint enrichment distinguish whether two genes or programs preferentially occupy the same ROI and whether their high expression occurs together. These measurements are calculated from expression at individual spots or cells, and from mean expression within tiles in each tessellation. Correlation measures whether the two expression values rise and fall together at the same spots, cells, tiles or hexagons. Bivariate Moran’s I (Wartenberg 1985; Anselin et al. 2002) instead relates one gene or program’s expression to the other’s expression in neighbouring units. Within each connected component, cross-correlation across distance bands describes how this relationship changes with spatial separation. At boundaries, TileScope compares expression near and far from a particular neighbouring ROI, separately for each feature and for each side of the boundary. This can reveal an expression pattern localized at the boundary that is not apparent from ROI-wide correlation.

Variation across tessellations quantifies sensitivity to the choice of partition, while comparisons between the original and aggregated representations reveal changes in effect magnitude or direction, making aspects of the MAUP explicit. Differences between representations are interpreted alongside their contributing observations, because aggregation can change weighting and coverage as well as spatial resolution. TileScope therefore reports availability and support alongside the measurements; analyses of the original observations can remain informative even when a component cannot be tessellated. The outputs are interpreted jointly with spatial maps, using the scenarios evaluated below to connect combinations of measurements to possible expression arrangements rather than assigning regions to mutually exclusive classes.

Together, these analyses describe where features are enriched, how they vary together within ROIs and their connected components, and whether their expression differs near boundaries with neighbouring ROIs, while assessing sensitivity to spatial partitioning and aggregation.

### TileScope distinguishes spatial configurations through complementary measurements

We designed a benchmark around biologically plausible arrangements of gene expression and tissue populations: localized program activity within a broader expression field, coordinated or opposing tissue transitions, embedded compartments, dispersed populations such as immune infiltrates, and expression changes at tissue boundaries. We used these constructions to test whether combinations of TileScope statistics provide recognizable evidence for particular spatial arrangements, beyond identifying positive or negative Pearson correlation. The benchmark comprised seven scenario families represented by ten modes, with three examples per mode (selected examples in Fig. 2A; complete catalogue in Supplementary Fig. S1). Controlled expression patterns were generated on observed tissue coordinates, retaining predefined ROI labels, disconnected components and contacts between regions (Supplementary Section 3).

**Figure 2.**
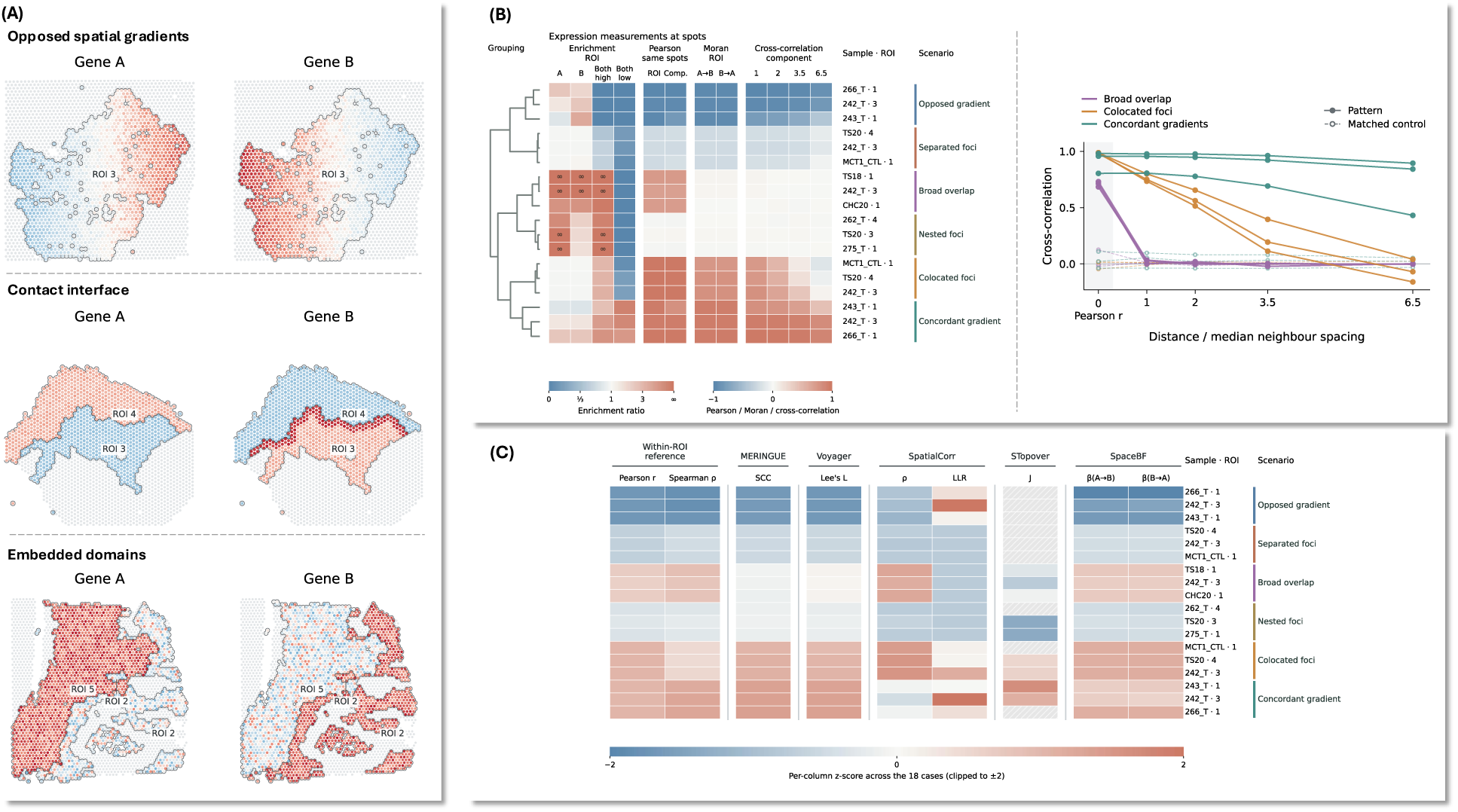
Complementary statistics distinguish simulated spatial configurations. **(A)** Simulated Gene A and B expression on observed tissue coordinates. Outlines mark ROIs; blue to red indicates increasing expression. **(B) Left**: 12 spot statistics for six modes, with three datasets per mode, clustered by average linkage after standardization and equal group weighting. Colours show enrichment ratios, Pearson correlations, bivariate Moran statistics and distance- band cross-correlations. **Right**: spot level Pearson correlation at distance zero and cross-correlation across four distance bands within each ROI’s largest component. Distances are relative to median nearest-neighbour spacing. Solid lines and filled symbols indicate patterns; dashed lines and open symbols indicate matched controls. **(C)** Comparator method statistics on the same 18 positive cases as (B) Left, grouped by method. Rows follow the same order. Each column shows the per-case value z-scored across the 18 cases and clipped to ±2. Hatched cells are out of a method’s scope. Method settings and statistic definitions are provided in Supplementary Section 4.5.

Clustering all 30 simulated datasets distinguished the six modes describing expression within ROIs, dispersed populations and the two boundary modes. For each of these nine modes, its three datasets formed a branch containing no other mode, without using scenario labels to determine the grouping (Supplementary Fig. S2; Supplementary Table 5). Embedded domains also formed a distinct group in an alternative analysis using Ward clustering restricted to statistics available in every dataset (Supplementary Table 6). Fig. 2B (left) illustrates the six expression modes using a smaller set of enrichment, Pearson correlation, bivariate Moran’s I and distance statistics.

Restricting clustering to ROI Pearson correlation and bivariate Moran statistics separated broad overlap, both focal modes and both gradient modes, but not the other five modes. The same five modes separated at spots, tessellations and direct and ensemble consensus representations (Methods Section; Supplementary Tables 5-6). Changing spatial representation therefore did not recover the additional distinctions obtained with the full statistical profiles.

In the nested foci scenarios, Gene B formed localized patches within an ROI where Gene A was broadly expressed. Gene B was enriched in the ROI, but the correlation between the two genes’ expression values was close to zero, showing that weak correlation did not exclude localized expression within shared territory. Dispersed populations also formed a distinct group despite their small B-labelled components lacking tessellation support; expression on these components remained available at individual spots.

Correlation alone mixed colocated foci with concordant gradients, whereas distance profiles alone mixed broad overlap with nested foci; the combined profiles separated both pairs. Spot cross-correlation illustrated the different spatial organization underlying positive expression correlation (Fig. 2B, right): broad overlap showed little correlation between distinct spots, colocated foci showed declining correlation with separation, and concordant gradients retained positive values across all four distance bands. Comparisons of scenario grouping at spots, tessellations and the two consensus representations are reported in Supplementary Table 6 and Supplementary Data 1.

In the contact interface scenarii that involve an anchor/reference ROI and a responder/target ROI2, although correlation between genes was weak across the responder ROI, boundary statistics distinguished increased Gene B expression beside an A-rich region (responder activation case) from Gene A expression extending across that region’s boundary (anchor’s extension case). Differences in mean expression between spots near and far from the boundary identified which gene carried the localized increase. Omitting these statistics prevented the two modes from forming separate branches (Supplementary Fig. S3; Supplementary Table 5).

To position these distinctions against existing spatial co-expression methods, we evaluated five published methods (MERINGUE, Lee’s L (Lee, 2001) implemented in Voyager (Moses et al. 2023), SpatialCorr, STopover and SpaceBF) and a non-spatial within-ROI Pearson and Spearman baseline reference on the same eighteen positive cases (Fig. 2C; Supplementary Section 4.5; Supplementary Data 1). The spatially weighted same-location statistics tracked the within-ROI Pearson reference closely, flipping sign between opposed and concordant gradients but mixing broad overlap with concordant gradients and separated foci with nested foci. SpatialCorr’s derived ρ separated broad cases from the focal modes; its likelihood ratio responded to the strongest effects without a directional sign. STopover quantified spatial overlap; separated foci and opposed gradients were excluded from this comparison (Supplementary Section 4.5). No single statistic from these methods distinguished the six expression modes on its own, consistent with the observation in Fig. 2B that correlation alone, or distance profiles alone, mixed pairs that only the combined profiles separated. Because these scenarios are deliberately clean, biological samples are expected to combine arrangements that are harder to tell apart than these; this evaluation therefore sets a “lower bound” on the statistical variety needed to characterise spatial expression relationships in observed tissue.

Together, these results provide a quantitative reference for recognizing the tested spatial arrangements from combinations of TileScope statistics. Such profiles can guide biological interpretation by distinguishing shared expression patches, extended transitions and expression concentrated near the boundary with a particular neighbouring ROI.

### Shared territories, internal covariation and boundary patterns distinguish glioblastoma program organization

Greenwald et al. (2024) characterised glioblastoma organization through program colocalization, adjacency and regional composition at multiple spatial scales, identifying a hypoxia-associated layered architecture. Building on this work, we examined how continuous program scores vary within individual ROIs and near their boundaries, separately from regional enrichment. Using their 14 transcriptional programs, we analysed 91 pairwise combinations across 192 BayesSpace ROIs in 30 *IDH1* wild-type (WT) glioblastoma sections: 12 profiled by Greenwald et al. (2024) and 18 by Ravi et al. (2022) (Supplementary Materials Section 5; Supplementary Data 2).

Across ROIs, selected neural/glial and reactive myeloid program pairs frequently showed positive Pearson correlations, whereas selected pairs involving the hypoxia-associated mesenchymal program (MES-Hyp) more often showed negative correlations (Fig. 3A). In sample 259_T, ROI 2 was enriched for astrocytic, macrophage and reactive-astrocytic programs but depleted for MES-Hyp. Within this ROI, reactive astrocyte program scores covaried positively with astrocyte-like malignant cell program (AC) and macrophage program (Mac, a signature that examines the inflammatory macrophages/myeloid cells) and negatively with MES-Hyp (Fig. 3B). These relationships were modest at individual spots (Pearson r = +0.14, +0.11 and −0.12, respectively) and stronger in the consensus representation (+0.56, +0.49 and −0.43), illustrating their dependence on spatial aggregation.

**Figure 3.**
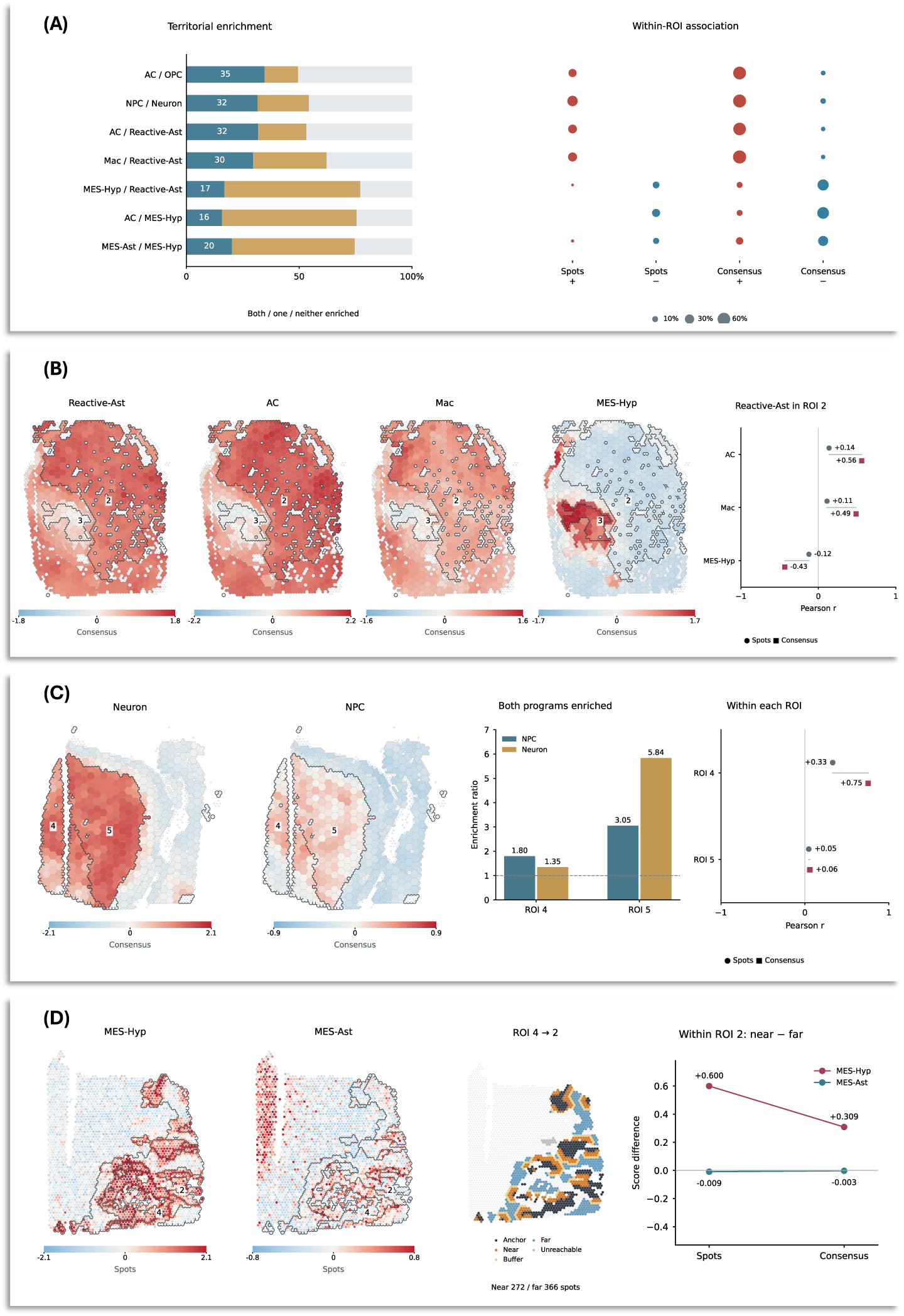
Regional enrichment, within-ROI correlation and boundary expression in glioblastoma. **(A)** Seven program pairs across 30 GBM samples. Bars show frequencies of both, one or neither program enriched relative to remaining tissue; numbers give percentages with both enriched. Bubble areas show frequencies of Pearson (r > 0.20) (red) or (r < -0.20) (blue). Frequencies use evaluable ROIs within samples, averaged with equal sample weights. **(B)** Sample 259_T consensus maps and correlations of Reactive-Ast with AC, Mac (inflammatory macrophages/myeloid cells) and MES-Hyp in ROI 2. Circles indicate spots; squares - consensus. **(C)** NPC and Neuron consensus maps, enrichment and correlations in ROIs 4 and 5 of sample 262_T. **(D)** MES-Hyp and MES-Ast spot maps, boundary bands and mean near-minus-far scores within ROI 2 of sample 260_T, relative to contact with ROI 4. Near and far groups are 0–1 and ≥ 3 neighbour steps from contact, respectively. Outlines mark ROIs; blue to red indicates increasing program scores. Consensus combines ten tessellations.

Regional enrichment and internal covariation nevertheless described different properties. In 262_T, neural progenitor-like malignant-cell (NPC) and neuronal (Neuron) programs were enriched in both ROIs 4 and 5. Their scores covaried in ROI 4 (r = +0.33 at spots; +0.75 in consensus) but had near zero Pearson correlations in ROI 5 (+0.05; +0.06), despite stronger enrichment there (Fig. 3C). Thus, the same programs could preferentially occupy two ROIs while varying together within only one. This distinction refines the recurrent NPC-Neuron association reported by Greenwald et al., separating shared regional enrichment from coordinated expression inside that region.

Boundary measurements revealed another form of organization in peripheral regions. In 260_T ROI 2, correlations of MES-Hyp with mesenchymal/astrocytic malignant cell program (MES-Ast) were near zero at both spots (−0.03) and consensus (−0.05) levels. Yet, within ROI 2 MES-Hyp scores were higher near its contact with ROI 4 than farther from that boundary, with near versus far differences of +0.60 and +0.31 at the two supports, respectively. MES-Ast showed almost no corresponding difference (Fig. 3D). The boundary measurement therefore identified a program-specific spatial pattern that the ROI-wide pair correlation did not describe.

Together, these examples show how TileScope complements recurrent program associations by distinguishing shared enrichment, internal covariation and expression concentrated near a particular boundary. These properties need not coincide, and their separation provides a more specific description of the organization within individual tissue compartments. Complete cohort measurements, support- sensitivity checks and additional spatial examples are provided in the Supplementary Material.

### Control and MCT1-knockout samples differ in tumour margin organization

To investigate the role of MCT1 in tumour-associated macrophages, we generated *Mct1^ΔMye^*mice by crossing *Mct1*^flox/flox^ mice with the LysM-Cre deleter strain (Cre recombinase driven by the *Lyz2* promoter predominantly expressed in myeloid cells). We generated 10x Genomics Visium profiles from orthotopic gliomas from *Mct1^flox/flox^* controls (CTL) or *Mct1^ΔMye^* mice (KO), together with an additional orthotopic glioma sample from a C57BL/6N WT mouse (WT-V4) (Supplementary Section 7.1). We examined how tumour-associated and immune-associated expression programs are organised within tumour regions and at their margins. The selected CTL and KO peripheries contained both proliferation and myeloid/phagocytic program expression, but the two programs showed contrasting spatial relationships. Common small gene signatures were scored per spot, and TileScope analyzed every available signature pair across all ROIs (222 sample-pair analyses; Supplementary Section 7; Supplementary Table 9; Supplementary Data 4).

We compared CTL periphery ROI 5 beside core ROI 2 with KO periphery ROI 9 beside core ROI 4, comprising 208 and 412 spots, respectively (Fig. 4A). Both programs were enriched relative to the other ROIs of each section. Their scores covaried positively within the CTL periphery (Pearson r = +0.20 at spots and +0.48 in ensemble consensus), but negatively within the KO periphery (−0.40 and −0.65; Fig. 4B–C). Both peripheries contained an excess of spots with high proliferation and myeloid scores relative to the other ROIs (7.01-fold in CTL; 4.60-fold in KO), but the two scores varied together in CTL and in opposite directions in KO.

**Figure 4.**
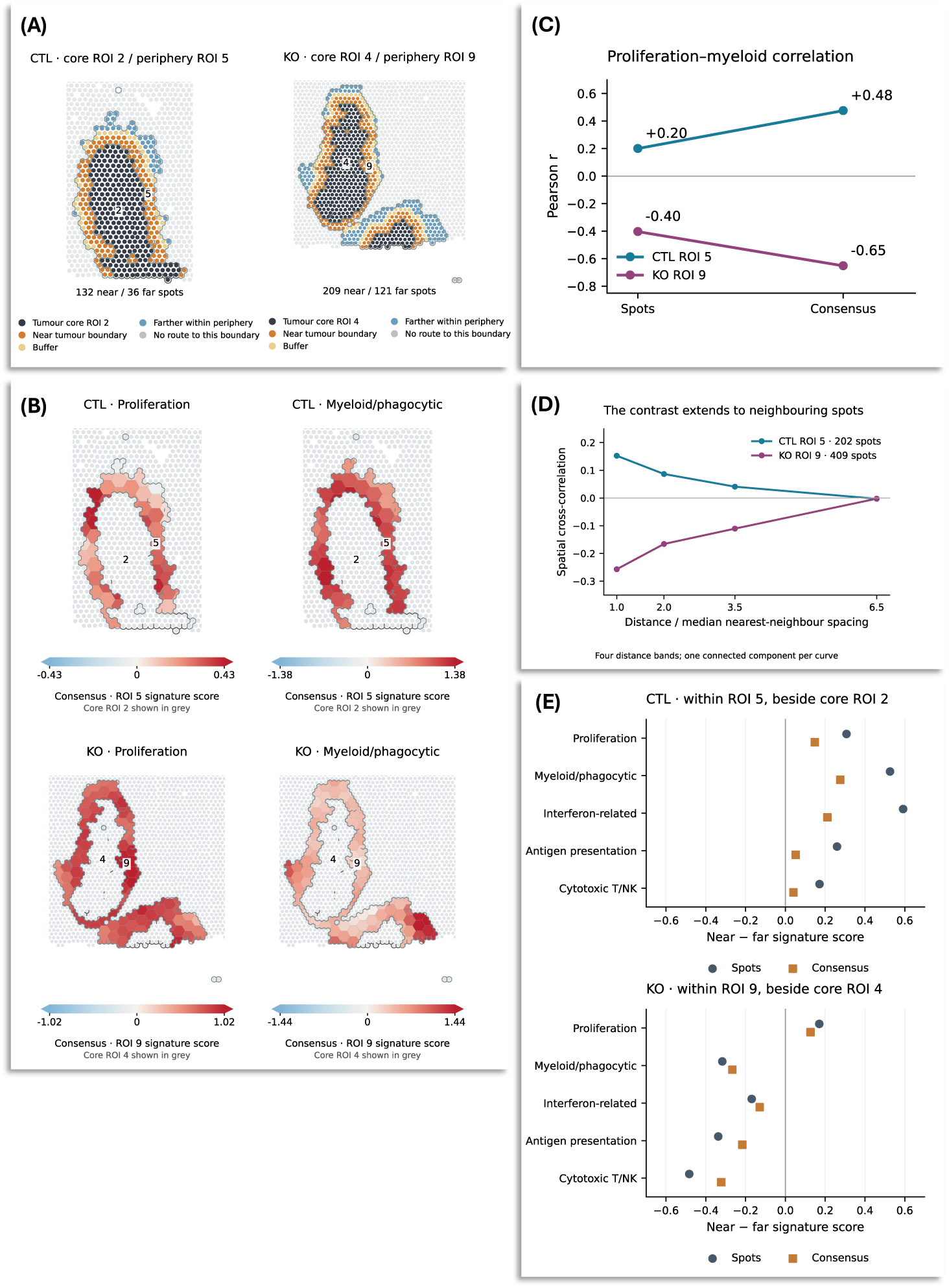
Contrasting organization of proliferative and immune-associated expression at control and MCT1 knockout tumour margins. **(A)** Tumour core and tumor periphery ROIs in CTL (ROIs 2 and 5) and KO (ROIs 4 and 9), with near tumour-facing boundary, buffer, farther within periphery and unreachable spots marked within each peripheral ROI. **(B)** Consensus maps of proliferation and myeloid/phagocytic scores; tumour cores are depicted in grey and ROI boundaries are outlined. Colour scales are set separately for each map. Corresponding spot maps are shown in Supplementary Fig. S9A. **(C)** Pearson correlation of proliferation and myeloid programs across periphery spots (CTL, 208; KO, 412) and consensus hexagons (58 and 110, respectively). **(D)** Spatial cross-correlation between these programs across four distance bands within the largest connected component of each periphery (202 CTL spots; 409 KO spots). Distances are scaled by each component’s median nearest-neighbour spacing. **(E)** Mean program score near the tumour boundary minus the mean farther within the same periphery, calculated at spots (circles) and in consensus (squares). Positive values indicate higher scores near the tumor-facing boundary. Consensus combines ten tessellations. Numerical results and analysis settings are provided in Supplementary Data 4.

Boundary measurements located this contrast relative to the tumour core. Within each peripheral ROI, we compared expression near its tumour-facing boundary (the contact zone with the ROI annotated as tumour core) with expression farther from that boundary into the same periphery. Proliferation scores were higher near the core-facing boundary than farther within the periphery in both sections. Myeloid scores were also higher near that boundary in CTL but were higher farther from it in KO (Fig. 4E). The proliferation-rich and myeloid-rich parts of the KO margin therefore occupied different positions relative to the core. These comparisons were made entirely within each peripheral zone.

The other immune-associated programs supported the same anatomical distinction. Interferon-related expression was positively correlated with proliferation in CTL and negatively correlated in KO and followed the myeloid near versus far contrast (Supplementary Fig. S9B; Fig. 4E). Antigen presentation and cytotoxic T/NK-cell scores remained positively correlated with each other in both peripheries but were higher near the tumour-facing boundary in CTL and farther from it in KO. These two immune-associated programs therefore remained positively associated in both sections, while their position relative to the core and to proliferation differed. Within the KO periphery, myeloid scores also covaried strongly with mesenchymal/stromal scores (r = +0.63 at spots and +0.91 in consensus), and both were higher farther from the core boundary (Supplementary Table 10; Supplementary Data 4).

Using the unweighted mean expression of each signature’s genes gave the same contrast: positive correlation in CTL (+0.45) and negative correlation in KO (−0.44). Correlations across ten alternative tessellations had medians of +0.47 in CTL and −0.54 in KO, with interquartile ranges of +0.32 to +0.48 and −0.59 to −0.52, respectively (Supplementary Fig. S9C). Nearby-spot cross-correlation was also positive in CTL and negative in KO, weakening towards zero at greater distances within each periphery’s largest connected component (Fig. 4D). The contrasting directions were therefore retained after aggregation and in the central ranges of the sampled partitions, while the distance profiles showed that the relationship was strongest locally.

Component-specific analysis revealed a further distinction within KO tumour ROI 4. A separate 49-spot component had mean tumour-associated and proliferation scores similar to those in the main 243-spot component (0.80 versus 0.79 and 0.89 versus 0.90, respectively). However, its correlation between interferon and proliferation programs was more strongly negative (r = −0.45 versus −0.09). Its cross- correlation was negative at short distances but positive in the most distant band, whereas the main component remained near zero (Supplementary Fig. S9D). Similar mean scores therefore concealed different internal expression relationships. The detached component’s location and tumour-associated expression identify it as a possible tumour satellite.

We also analyzed a tumour sample from a WT C57BL/6N mouse lacking the floxed allele and Cre transgene (sample WT-V4). The WT-V4 sample provided a complementary example of expression organized at a tumour boundary. In the computationally annotated tumour-brain interface (ROI 2) reactive-interface and myeloid scores covaried positively (r = +0.48 at spots and +0.81 in consensus). Both were higher near tumour core ROI 10, and positive cross-correlation extended across the measured distance bands within the largest connected component. A common mesenchymal/stromal signature recovered a related pattern (+0.57 and +0.81), showing that the observation was not confined to the WT-V4-specific reactive signature (Supplementary Fig. S10).

Together, these analyses distinguish the presence of expression programs from their organization around the tumour. Regional enrichment identified proliferative and immune-associated expression in both CTL and KO margins; internal correlations and boundary measurements showed how their positions differed. Keeping disconnected components separate revealed another local relationship within a single tumour label. TileScope linked these findings to the regions, components and boundaries where they occurred. The comparison of CTL and KO samples motivates testing whether MCT1-dependent tumour–myeloid interactions contribute to margin organization, lymphocyte infiltration and anti-tumor immunity.

## DISCUSSION

TileScope distinguishes spatial scenarios that can produce similar enrichment or correlation values. It combines measurements of shared regional expression, covariation within regions, cross-correlation over distance and expression near specific neighbours. Each measurement remains linked to the tissue region, connected component or boundary on which it was calculated. This allows different relationships between the same programs to be examined within one sample, including patterns confined to a small component or a particular contact. Repeated partitioning adds an assessment of how these measurements change when spots or cells are grouped differently.

No individual statistic evaluated in the benchmark described all the tested spatial scenarios. Similar spot level correlations occurred in coincident patches and aligned gradients, while expression concentrated near a boundary could accompany weak correlation across the region. Distance profiles and boundary measurements supplied information that resolved several of these ambiguities. The constructions used controlled continuous signals on tissue geometries observed in experimental data, allowing the effects of particular spatial changes to be examined without reproducing the full variability of measured expression. Several statistical families supplied overlapping information, and separation of embedded domain cases depended on the statistical subset and clustering method. These scenarios provide reference profiles for interpreting measurements alongside expression maps. They do not determine a unique geometry from a statistical profile, and real tissue may combine several arrangements.

The external-method comparison further illustrates the different information supplied by spatial cross- correlation, spatially varying correlation, local regression and component overlap. These are the respective targets of approaches such as MERINGUE, SpatialCorr, SpaceBF and STopover (Miller et al. 2021; Bernstein et al. 2022; Seal and Neelon 2026; Bae et al. 2025). None of the individual summaries evaluated distinguished all six within-ROI expression modes. The comparison supports combining measurements of different spatial properties, without establishing a general ranking of the complete methods.

The human glioblastoma analysis shows what these distinctions add to recurrent program relationships described across samples and spatial scales (Greenwald et al. 2024). NPC and neuronal programs were enriched in two ROIs but correlated within only one. Hypoxia-associated expression also increased near a specific neighbour despite negligible pairwise correlation across the responder ROI. A published program relationship thus leaves open whether the programs share tissue preference, vary together within a compartment or exhibit a local boundary pattern. Locating these differences identifies specific tissue contexts for biological follow-up.

In our own novel orthotopic mouse glioma datasets, the combined measurements gave anatomical meaning to a contrast between control and myeloid-targeted MCT1-knockout sections. Proliferation and myeloid programs were enriched in both selected peripheries, yet their scores were positively correlated in control and negatively correlated in knockout. Proliferation was higher near the core-facing boundary in both samples, while myeloid expression was higher near that boundary in control and farther within the knockout periphery. The LysM-Cre manipulation targets myeloid populations, including macrophages and granulocytes (Clausen et al. 1999). Visium cannot assign the observed differences to individual recombined cells, but it locates the tissue in which their consequences might be investigated. The observation motivates testing whether myeloid MCT1 influences the positioning and expression states of cells at the tumour margin.

These applications demonstrate the continuing value of spot-based datasets for questions about tissue organization. Regions already annotated by cell state or enrichment can be revisited to examine their internal variation, disconnected components and relationships to neighbouring compartments. The resulting spatial descriptions can guide cellular measurements or experimental sampling even when the original assay cannot resolve individual cells.

Averaging nearby measurements helps describe expression patterns across groups of spots or cells, but requires choosing which observations belong together. Even within a well-defined tissue zone, different averaging units can produce different estimates of the relationship between programs. TileScope assesses this dependence by repeatedly partitioning the same ROI and recalculating the statistics, addressing an aspect of the modifiable areal unit problem (MAUP; Stillwell et al. 2018). In the mouse analysis, positive correlation between proliferation and myeloid programs in control and negative correlation in knockout persisted in the central ranges of the tessellation estimates, reinforcing this observation. Selected human glioblastoma correlations became substantially stronger after aggregation. These comparisons make the averaging choices part of the interpretation. A conclusion such as coordinated expression within a tumour margin can be assessed across alternative partitions, showing how much it depends on the placement of boundaries. Comparisons with the original observations also establish whether that conclusion holds at spot or cell resolution.

We applied TileScope to expression-based ROIs in Visium data and to cellular neighbourhoods and pathological compartments in Xenium data, extending the analysis to large datasets of irregularly spaced cells. The Xenium glioblastoma regions were defined from local cell type composition, while the breast regions were delineated with histopathological and single cell transcriptomic information (Meylan et al. 2026; Janesick et al. 2023). Selected expression contrasts reported in these studies remained apparent after aggregation, demonstrating the use of TileScope beyond regions obtained by clustering expression profiles. Notice that we developed the scenario catalogue with expression-defined regions in mind. Anatomical annotations can encompass several transcriptionally distinct compartments, so differences within an anatomical ROI may reflect the organization of those compartments. Applying the scenarios in this setting may require adapting their descriptions to the meaning and granularity of the supplied regions.

Further development can address arrangements that remain incompletely described, particularly expression within individual labelled inclusions whose parent ROI also occupies other parts of the section. The scenario catalogue supplies controlled comparisons for assessing what an additional measurement captures, its overlap with existing statistics and its remaining ambiguities. TileScope provides a reproducible way to examine gene and program relationships within annotated tissue compartments, identify the components or boundaries carrying a pattern, and assess its dependence on spatial grouping.

## METHODS

### Analysis design and input data

TileScope describes relationships between two genes or expression programs within predefined tissue regions and at their boundaries. It distinguishes regional enrichment, within-ROI Pearson correlation, bivariate Moran’s I, cross-correlation across distance bands, and expression differences near a specified neighbouring ROI. ROI labels remain fixed while repeated spatial partitions provide alternative groupings of the same spots or cells. Statistics retain their ROI, component or boundary identity, allowing different spatial arrangements to coexist within a sample rather than assigning the entire sample one relationship.

Each sample supplies planar coordinates, expression values values *x_i_*, *y_i_* and an ROI label for every spot or cell *i* ∈ *V*. An ROI *R* ⊆ *V* contains spots or cells sharing a label and may have disconnected components. Distances are expressed relative to sampling spacing. Spot- or cell-level expression remains available even when the corresponding components cannot support spatial aggregation.

### Spatial connectivity and repeated tessellation

A sample-wide undirected graph *G* = (*V*, *E*) represents spatial neighbourhoods. For each spot or cell *i*, *d_i_* is the distance to its nearest other spot or cell in the same sample; *d*^/^ is the median of these distances across the sample. Visium edges connect spots separated by at most *τ_V_d*^/^. Xenium Delaunay edges of length *P_i_*_j_ are retained when both *P_i_*_j_ ≤ *τ*_local_max(*d_i_*, *d*_j_) and *P_i_*_j_ ≤ *τ*_cap_*d*^/^. The dimensionless distance multipliers

are specified in Supplementary Table 2. Within an ROI, connected components are computed from the induced graph *G*[*R*], while edges between different ROI labels identify contacts. Graph construction is independent of the expression pair.

Voronoi tessellation partitions the area of each connected component *C* into spatial tiles within which expression is averaged. To delimit the area for tessellation, we first construct a polygon for each *C*, combining small discs centered on each spot or cell with Delaunay triangles whose three edges belong to *G*[*C*]. This area is clipped using the sample coordinates and ROI labels to limit extension into unsampled areas or neighbouring ROIs; discs around spots or cells excluded from tessellation are removed (Supplementary Section 1). Each separate polygon containing enough spots or cells to construct a tile is processed independently. We denote this polygon by *D* and refer to it as a tessellation domain.

For each tessellation domain *D*, TileScope generates *K* Voronoi partitions T^(1)^(*D*), … , T^(*K*)^(*D*). Seeds are sampled without replacement from its *N_D_* unique observation coordinates and slightly perturbed. For a target of *q* spots or cells per tile, the initial number of seeds is

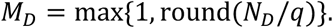

Tiles containing fewer than the specified minimum number of spots or cells, or having an area below a geometry-based threshold, are merged with neighbouring tiles, preferentially along the longest shared boundary.

For tile *t* in partition *k*, let 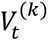 contain its assigned locations and 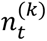 their number. Tile expression is their mean,

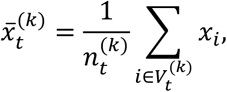

with 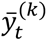 defined in the same way. At each partition index, ROI statistics pool tiles from all retained polygons of that ROI. Repeated partitions assess sensitivity to the spatial grouping of expression values, following the use of alternative partitions to investigate the modifiable areal unit problem (MAUP; Stillwell et al., 2018).

### Ensemble consensus and direct aggregation

Within each connected component, a common hexagonal grid combines the different tessellations. Hexagons are clipped separately to each tessellation domain *D* of that component. Their size is determined from tile areas and the density of spots or cells, subject to an upper area limit (Supplementary Section 1.3).

Ensemble reconstruction uses the principle of areal interpolation (Goodchild and Lam, 1980). The overlap weight

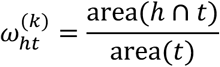

transfers the same fraction of tile *t*’s expression sum and spot or cell count to hexagon ℎ. The ensemble estimate is

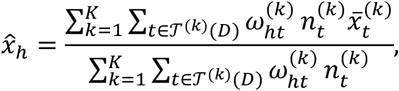

and likewise for *y*. This assumes uniform redistribution within each tile and weights partition contributions by the spot or cell counts projected onto each hexagon. High- and low-expression counts are transferred with the same weights. Hexagons must meet geometric overlap and tessellation-coverage requirements. Thus, the occupancy parameter controls grid sizing (Supplementary Section 1.3).

For comparison, direct consensus takes the mean expression of spots or cells assigned to each retained hexagon, without projection from tiles. Direct and ensemble estimates therefore share grid geometry but not necessarily the same contributing observations: a tile overlapping ℎ can project contributions from

observations outside ℎ. Direct estimates bypass this projection step. Empty or inadequately supported estimates are reported as unavailable.

### ROI enrichment

TileScope computes how frequently high or low expression occurs in an ROI, and whether expression levels are generally higher or lower than in the other ROIs.

Sample-wide expression quantiles define high and low values for each gene or program and are fixed across ROIs. A criterion *H* denotes high *x*, high *y*, joint-high or joint-low expression; joint criteria require both values to satisfy their threshold. With *R̅* denoting spots or cells in the other labelled ROIs, enrichment of *R* is

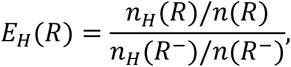

where *n_H_* counts spots or cells satisfying *H* and *n* is their total number in *R* or *R*^–^, irrespective of whether they satisfy *H*. Values > 1 indicate greater prevalence than in the reference. Just as individual enrichment, joint enrichment compares joint occurrence with the other ROIs.

Tessellation enrichment counts tiles according to whether their mean expression satisfies *H*, separately for each tessellation. For ensemble consensus enrichment, each tile’s total spot or cell count and its count satisfying *H*, are distributed onto the consensus hexagons according to tile–hexagon overlap. These counts are averaged across contributing tessellations, then used to calculate enrichment in the ROI relative to the other ROIs.

TileScope also compares expression levels within the ROI and in the other ROIs. It reports the probability that an expression value from the ROI exceeds one from the reference, and a log2 ratio of their median or outlier-resistant mean expression, selected according to sample size (Supplementary Section 2.1).

### Correlation and bivariate Moran’s I

Within each ROI, Pearson correlation between the two genes or programs is calculated using spot or cell values, tile mean expression values separately for each tessellation, as well as for direct and ensemble consensus values. Values from contributing components are combined in each ROI calculation, so correlation can reflect both variation within components and differences between their mean expression levels. Correlations obtained from repeated tessellations are summarized by their median and interquartile range.

Bivariate Moran statistics compare one gene expression with neighbouring values of the other. For any one representation, let *u_i_*, *v_i_* be the expression values standardized within the ROI, *n* the number of finite paired units, *W* = (*w_i_*_j_) the spatial weights and *S*_O_ = ∑*_i_*_,j_ *w_i_*_j_. Then

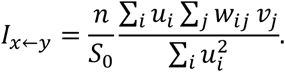

Exchanging the genes gives *I_y_*_➛_*_x_*. For spots or cells, weights are obtained by row-standardizing *G*[*R*]. For tiles and hexagons, mutual nearest-neighbour graphs of unit centers are constructed within each retained polygon, combined without between-polygon edges and row-standardized. The two directions identify which gene supplies neighbouring values.

Constant expression, insufficient neighbouring units or inadequate coverage yield unavailable results. For tiles or hexagons, Moran’s I is unavailable when, within every tessellation domain *D*, each unit is connected to all the others, leaving no distinction between nearby and distant neighbours. Eligibility is assessed separately for each statistic (Supplementary Section 2.3).

### Cross-correlation over distance

Distance profiles use expression within each retained connected component *C*. Euclidean distances are divided by the component’s median nearest-neighbour spacing and assigned to fixed bands. Expression values 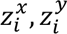 are standardized within *C* to zero mean and unit population variance. For a band *b*, let N*_b_*(*i*) contain neighbours of spot or cell *i* in that band, and A*_b_* the spots or cells with at least one such neighbour. The symmetric cross-correlation is

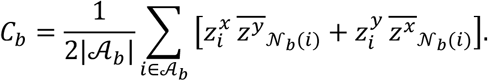

Bars denote means over the indicated neighbours. Each contributing spot or cell has equal weight, regardless of its number of neighbours. Bands require a minimum contributing count and fraction. Expression is standardized once within the component. The resulting cross-correlations *C_b_* need not lie in [−1,1] as they are not Pearson coefficients.

### Expression near neighbouring ROIs

The boundary response is defined on an ordered pair of contacting ROIs. For contacting ROIs *Q* and *R*, the reference or “anchor” ROI *Q* identifies the contact, and the “responder” ROI *R* is the one in which the measurement is made: the response evaluates whether a selected feature is higher in *R* near its contact with

*Q* than farther inside *R*. “Responder” denotes the ROI in which the measurement is made; it does not imply that the expression change originates there. Contact spots or cells are those in *R* joined to *Q* by an edge of *G*. Distance from the contact is the shortest path within *G*[*R*]; spots or cells that cannot be reached from the contact are omitted. Near and far groups are separated by an excluded buffer. For a gene or program *x*, the boundary response is calculated from individual spots or cells within *R* as

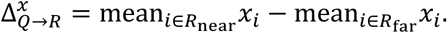

Both means concern the responder. Anchor’s ROI label identifies the contact, but its expression does not enter the calculation. The response is calculated for each gene and in the reverse ROI order. Near and far groups are defined using individual spots or cells. These assignments are carried into tile and hexagon calculations through the corresponding counts and overlap weights.

Tile and hexagon expression values are weighted by their respective near and far counts when calculating the two group means. Minimum group sizes and coverage requirements are specified in Supplementary Table 2.

### Sensitivity to partitioning and aggregation

The median, interquartile range and number of valid tessellations summarize dependence on alternative partitions at the chosen target occupancy. Comparing the same statistic calculated at individual spots or cells, across tessellations and on the consensus grids identifies changes in direction or magnitude after aggregation. Enrichment comparisons use the transformed ratio *E_H_*(*R*)/j1 + *E_H_*(*R*)k − 1/2.

Partition variability and differences between representations address different aspects of MAUP. The former changes observation groupings within the same aggregation procedure; the latter can change resolution, weighting, represented observations and, for enrichment, the counting unit.

For each ROI and statistic, TileScope reports whether the effect retains its sign across spots or cells, tessellations and ensemble consensus, varies substantially between tessellations, or reverses sign after aggregation. A sign reversal means that Pearson correlation or bivariate Moran’s I changes from positive to negative, or conversely. For enrichment, it indicates a switch between enrichment and depletion relative to the other ROIs. These interpretations consider effect magnitude and variation between tessellations; weak effects and unavailable comparisons are reported separately. The criteria are specified in Supplementary Section 2.4.

### Software availability

TileScope is open-source and available at https://github.com/cbib/TileScope. The scripts used to generate the results in this paper are in the companion repository https://github.com/cbib/TileScope-results .

## DATA ACCESS

Raw and processed sequencing data generated in this study have been submitted to the NCBI Gene Expression Omnibus (GEO; https://www.ncbi.nlm.nih.gov/geo/) under accession number GSE351684

The companion datasets necessary to reproduce the results and intermediate analysis of the present publication are available via Zenodo (https://zenodo.org/records/23535235).

## ETHICS STATEMENT

All mice were housed under specific pathogen-free conditions at the Animalerie Mutualisee de Talence (University of Bordeaux, France) on a 12-h dark/light cycle with unlimited access to food and water. All experiments were performed with 8- to 11-week-old male mice and were randomly allocated to the treatment groups. Black6 mice were born, housed, and treated in the same animal facility. All animal procedures have been done according to the institutional guidelines and approved by the local ethics committee (agreement number #27715).

## ACKNOWLEDGEMENTS

The authors thank the French National Cancer Institute (INCa) for financial support through its PLBIO program (grant no. PLBIO2022-027).

## AUTHOR CONTRIBUTIONS

MAL: Conceptualization, Data curation, Formal analysis, Investigation, Methodology, Software, Validation, Writing: review and editing

SL: Conceptualization, Investigation, Orthotopic mouse experiments, Data interpretation, Writing: review and editing

BD: Data curation, Software

DCF: Initial data analyses, Writing: review and editing

MS: Funding acquisition, Experiment design, Writing: review and editing

TD: Conceptualization, Investigation, Validation, Funding acquisition, Resources, Writing: review and editing

MN: Conceptualization, Formal analysis, Funding acquisition, Investigation, Methodology, Software, Supervision, Writing: original draft

